# A polarized nucleus-cytoskeleton-ECM connection controls collective migration and cardioblasts number in *Drosophila*

**DOI:** 10.1101/2019.12.17.879932

**Authors:** C Dondi, B Bertin, JP Da Ponte, I Wojtowicz, K Jagla, G Junion

## Abstract

The formation of the cardiac tube is a remarkable example of complex morphogenetic process conserved from invertebrates to humans. It involves coordinated collective migration of contralateral rows of cardiac cells. The molecular processes underlying the specification of cardioblasts prior to migration are well established and significant advances have been made in understanding the process of lumen formation. However, the mechanisms of collective cardiac cells migration remain elusive. Here we identified *CAP* and *MSP-300* as novel actors involved in cardioblast migration. They both display highly similar temporal and spatial expression pattern in migrating cardiac cells and are required for the correct number, alignment and coordinated directional migration of cardioblasts. Our data suggest that CAP and MSP-300 are part of a tension-sensitive protein complex linking cardioblast focal adhesion sites to nuclei via actin cytoskeleton and triggering coordinated cardioblast movements.

## Introduction

The genetic network regulating heart development is highly conserved from flies to humans with orthologous transcription factors (TFs) families and signaling pathways essential for heart development such as Tinman/Nkx2.5, Pannier/GATA4-6, Tailup/Isl1, Dorsocross H15 Mid/Tbx, Hand, Dpp/BMP or Wg/Wnt ^1–5^. However, while mechanistic insights have been gained on the specification of cardiac progenitors, less is known about genes controlling subsequent events of cardiogenesis.

Just as in *Drosophila*, formation of the mammalian heart starts by the migration of bilateral strips of cardiac progenitor cells to the midline and assembly of linear heart tube. *Drosophila* provides thus an accurate model system to study early steps of cardiac morphogenesis.

Cardioblasts (CBs) migration occurs concurrently with dorsal closure a process leading to the spreading of the epidermis dorsally over an extraembryonic epithelium, the amnioserosa. Migrating CBs extend projections from their dorsal surface and towards the leading edge of the epidermis suggesting coupling of these two tissues^6,7^. However, during late phase of migration CBs move cell autonomously. They send filopodia that contact contralateral CBs and ensure correct linear heart tube assembly^6,8^.

A few actors involved in this process have been identified including the transmembrane protein Toll^9^ and participating in adhesion between ipsilateral CBs faint sausage^10^.

Proteins involved in setting or maintaining polarity of CBs have also been implicated in collective migration. Among them the Integrin dimer expressed by the CBs (αPS3, βPS), its ligand Laminin-A or cytoplasmic Integrin adapter Talin are all required for CBs alignment ^11–13^.

Moreover, the coordinated migration and alignment of CBs requires the Slit morphogen and its receptor Robo^14,15^. Slit functions in migration are conserved from *Drosophila* to vertebrates. In zebrafish, Slit2 knock down embryos show disruption of endocardial collective cell migration with individual cells migrating faster and with loss of directionality^16^.

Recent findings have shown a genetic interaction between αPS3 gene *scb*, and Slit/Robo suggesting an integrated function of all these cell membrane-associated actors to polarize CBs migration during cardiac tube formation^12^.

At a sub-membrane level, link between cell adhesion and actin cytoskeleton remodeling is also fundamental for CBs migration and requires intermediate proteins to mediate connections and transduce signals. Among them, two members of the Dock Guanine nucleotide Exchange Factors (GEFs) containing SH3 domain, Myoblast city and Sponge have been shown to be required during CBs migration through activation of Rac and Rap1, respectively. Mutants for these two genes revealed defects in the migrating CB rows with breaks and clusters of multilayered CBs^17^. Moreover, the Rho GTPase CDC42 is required in Tinman CBs for the contact with the opposite row^18,19^ by modulating nonmuscle myosin II Zipper localization^19^. These cytoskeleton modulations allow filopodia formation at the leading edge, tension sensing to keep CBs aligned, cell shape changes and might be transmitted directly to nuclei to coordinate their shape and functions^20^.

Here we identify two new actors of collective CB migration, Cbl Associated Protein (CAP) and Muscle Specific Protein 300 (MSP-300). CAP is thought to act at membrane-cytoskeletal interface of stretch-sensitive structures whereas MSP-300 provides a link between actin cytoskeleton and perinuclear layer *via* Linker of Nucleoskeleton and Cytoskeleton (LINC) complex and controls nuclei positioning in striated muscles. In the developing heart, our data show that CAP and MSP-300 accumulate in dots on apical and basal sides of migrating CBs and are both required for proper CBs alignment, Slit subcellular localization and proper CBs nuclei properties. We also demonstrate that polarized accumulation of nuclei-anchored MSP-300 and cell membrane associated CAP ensure cell cycle exit and directional CBs migration. These findings imply the requirement of nucleus-cytoskeleton-extracellular matrix connections during collective CBs migration.

## Results

### CAP and MSP-300 accumulate inter-dependently in apical and basal spots in migrating cardioblasts

To identify new actors of CBs migration we performed Tin-CB-specific TRAP-seq experiments (unpublished data) to capture engaged in translation transcripts during the course of heart closure (Sup Fig 1 A-C). Among identified candidates we first tested *CAP* and *MSP-300* as their functions have not yet been investigated in *Drosophila* heart while having orthologous genes, *SORBS* family and *Syne1* respectively, with conserved cardiac expression in mammals^21–24^.

*In vitro*, SORBS proteins are generally targeted to focal adhesion sites where they play diverse roles in mechano-transduction, contributing to stiffness sensing and contractility^25^. CAP is the only SORBS family member in *Drosophila* and has been described as an adaptor protein making links between different partners at membrane-cytoskeletal interface of stretch-sensitive structures *via* the vinculin-integrin complex^26^.

MSP-300 is a large protein of the Nesprin family with several protein domains including spectrin and nuclear anchoring KASH domains. MSP-300 has been described to participate in nuclei positioning in striated muscle fibers connecting actin cytoskeleton and perinuclear layer through the LINC complex.

Recently, MSP-300 has been shown to display stretching capacity participating in the elasticity of a myonuclear scaffold to protect myonuclei from mechanical stress^27^.

Nucleus deformability is necessary when cells are squeezing through small constrictions during their migration^28^. The role of the nucleus in cell migration is suggested to be dependent on its connections with components of the cytoskeleton mediated by LINC and indirectly with extracellular matrix including focal adhesion complex.

Because coordinated CBs migration towards the midline involves stretch induced morphological changes of cells we asked whether CAP and MSP-300 could be involved in these processes. We first tested expression patterns of CAP and MSP-300 by immunostaining and confirmed that they are both expressed in CBs during migration from late stage 13 (Fig. 1A-A”). Higher magnification pictures show that CAP and MSP-300 are co-expressed in a stereotyped dotty pattern on apical and basal sides of each CB clearly visible from stage 15 (Fig. 1B-B” and arrowheads in Fig. 1C-C”). This expression is maintained when the two rows of CBs have reached the dorsal midline (Fig. 1D-E”). While expression of CAP seems to be also diffuse under cell membranes (arrow in Fig 1E’), MSP-300 in contrary is very localized where the maximum of CAP protein is accumulated (arrowheads in Fig. 1 E”). Transversal sections of 3D reconstructed developing hearts confirmed co-localization of CAP and MSP-300 on basal and apical sides of CBs in aorta as well as in the heart proper (arrows in Fig. 1F-F”, Sup fig 1D-D’). To validate CAP and MSP-300 localization in the apical side at the level of the luminal domain we performed double immunostaining experiments with CAP and Slit antibodies (Sup Fig 1E-G”). We confirmed that CAP and MSP-300 are enriched apically at the level of the lumen (arrow in Sup Fig 1F’) but absent at the CBs adhesion sites characterized by Armadillo expression (Sup Fig. 1H-H’). Unexpectedly, we also observed Slit localization in a dotty pattern at the basal side of CBs at stage 15-16 similarly to CAP and MSP-300 expression pattern (arrowheads in Sup Fig. 1E-E”). This dotty accumulation of Slit is transient as it becomes polarized at the luminal side at late stage 16 (Sup Fig. 1G-G”).

**Figure 1:**
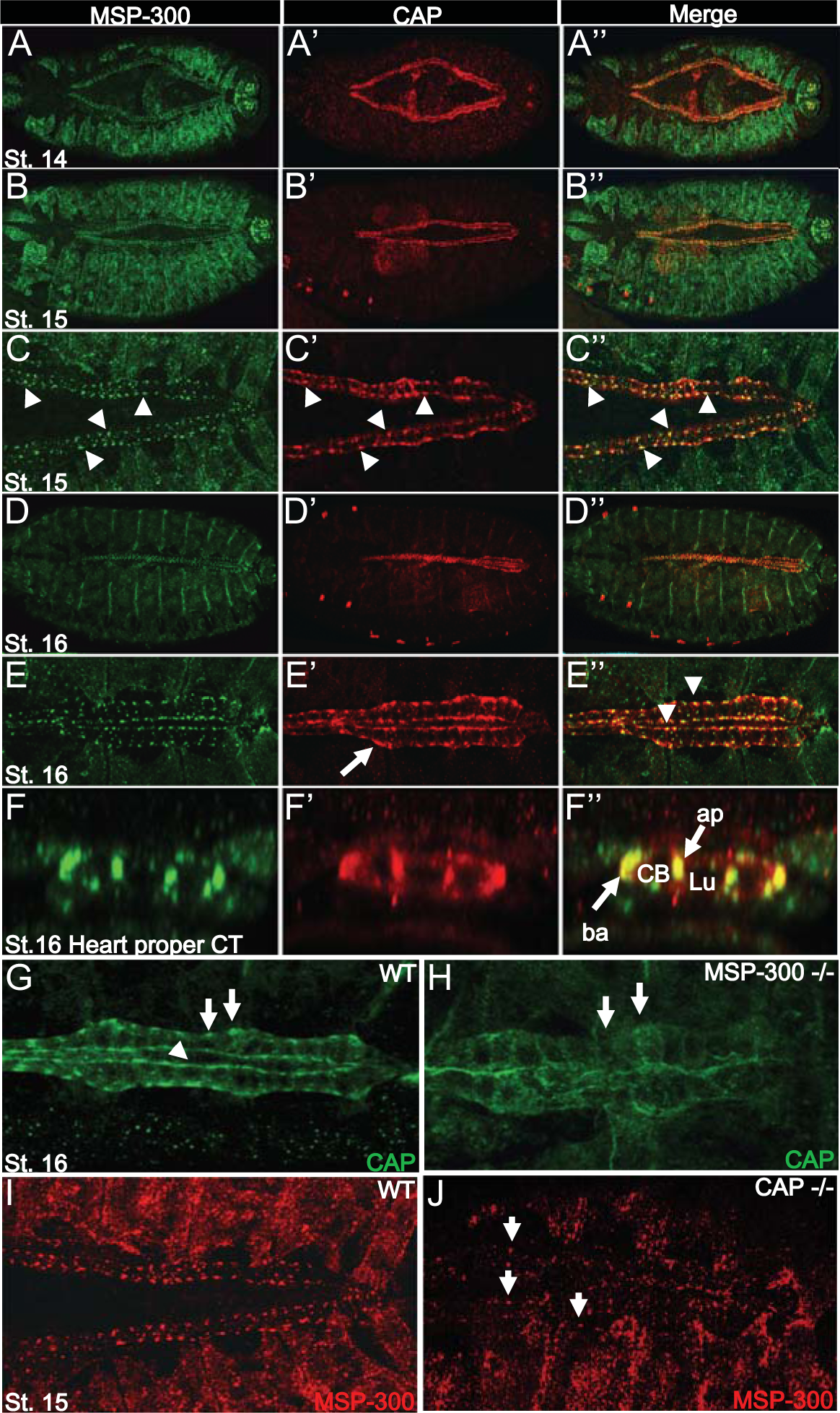
CAP and MSP-300 colocalize in CBs on basal and apical dots inter-dependently. (A-A”) Dorsal view of a stage 14 embryo showing by immunostaining with anti-CAP and anti-MSP-300 antibodies co-expression in cardioblasts. (B-B”) Dorsal view of an early stage 15 embryo showing that CAP and MSP-300 accumulate in dots on the basal and apical sides of cardioblasts. (C-C”) Higher magnification showing that MSP-300 accumulates where the maximum of CAP proteins on apical and basal sides (arrowheads) is present. (D-D”) Dorsal view of a stage 16 embryo showing that CAP and MSP-300 dotty pattern is present until the stage of lumen formation. We observe also a colocalization in muscle attachment sites at segmental borders. (E-E”) Higher magnification showing that CAP accumulation is extended under the cell membrane (arrow) and that CAP and MSP-300 colocalize on basal and luminal sides. (F-F’’) 3D reconstruction of transversal cut in heart proper validating accumulation of MSP-300 and CAP on basal and apical side of CBs (G-H) Dorsal view of a stage 16 embryo showing that loss of MSP-300 induces a strong decrease in CAP localization on basal (arrows) and apical sides (arrowhead). (I-J) Dorsal view of a stage 15 embryo showing that loss of CAP leads to a partial reduction of MSP-300 dotty pattern. Some dots are still visible in few CBs (arrows)

Next, we wondered whether CAP and MSP-300 are interdependent for their dotty localization. MSP-300 is considered as a scaffold protein with multiple domains so we hypothesized that it could participate in the recruitment and accumulation of CAP in subcellular compartment creating this repeated dotty pattern. Hence, we performed immunostainings in mutant contexts for these two genes. We observe a clear loss of dotty accumulation of CAP on basal and apical CB sides in *MSP-300^SZ75^* mutant embryos suggesting that MSP-300 is required for CAP localization in cardioblasts (arrows Fig 1G-H). Similarly, in *dCAP49e* loss of function context, MSP-300 pattern is severely affected. Because some of MSP-300 dots in CBs persist, we conclude that CAP is only partially required for MSP-300 spotty pattern (arrows Fig 1 I-J).

In summary, we observed that during the late phase of CBs migration, CAP, MSP-300 and Slit co-localize in polarized dots on each side of CBs in basal and luminal domains. As this spotty localization appears interdependent it suggests that CAP and MSP-300 are associated in interconnected units.

### MSP-300 and CAP are required for the correct alignment and number of Tin cardioblasts

We next examined *CAP* and *MSP-300* functions during CBs migration. Immunostaining with Mef2 antibodies revealed misalignments of CBs in both *MSP-300* and *CAP* mutant contexts (Fig. 2A-D”). Cardioblasts formed clusters (arrowheads in Fig. 2B,C) and multiple layers of cells (brackets Fig. 2B,C). We scored these alignment defects and found that 80% of *MSP-300* and 65% of *CAP* mutant embryos present a multilayered row of CBs in a portion of segments thus suggesting that *MSP-300* and *CAP* are both required to maintain cohesion of migrating CBs.

**Figure 2:**
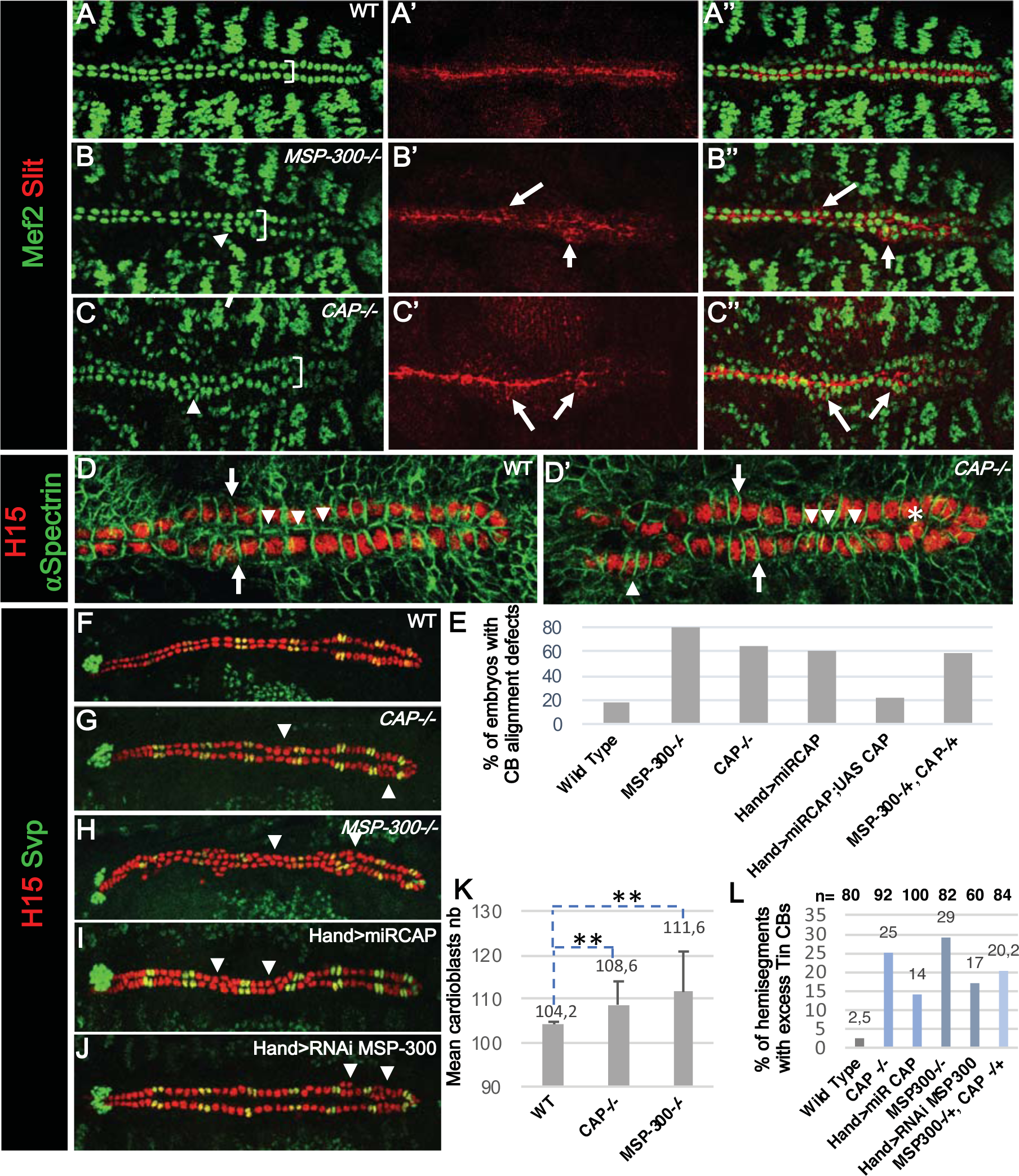
MSP-300 and CAP are required for correct CBs alignment and number during migration. (A-C’’) Dorsal view of stage 16 embryos showing that compared to WT, the loss of MSP-300 and CAP induces alignment defects visible with CBs nuclear marker Mef2 and CB polarity marker Slit. A-A” show the WT context with 2 rows of CBs (bracket) and apically localized Slit protein. B-B” In loss of *MSP-300* more than 2 rows of CBs are visible in some segments (bracket) and Slit pattern is not polarized with some accumulation visible on lateral and basal side of CBs. C-C” *CAP49e* mutant embryos show similar phenotypes than the loss of *MSP-300*. (D-D’) Cell membrane marker α-Spectrin shows that in *CAP49e* mutant cells are misaligned, and matching between contralateral CBs is disturbed (compare arrows in D and D’) impacting contacts between CBs (compare arrowheads in D and D’). Mislocalized cells are also visible in *CAP* mutant (star). (E) Histogram showing the percentage of embryos with CB alignment defects (n=30). (F-J) Dorsal view of stage 16 embryos revealing specifically with anti-H15 antibody all cardiomyocytes including Svp ostia cells. *CAP49e* and *MSP-300sz75* homozygous mutants as well as cardiac specific knock down for these two genes induce an increase in Tin CBs number in some hemisegments while Svp CBs number remain unchanged. (K) Histogram showing quantification of the mean CBs number. In *dCAP49e* (108,6) and *MSP-300sz75* (111,6) CB number is increased compared to the WT (104) (n=30 Student’s T-Test pval< 0,01). (L) Histogram showing the percentage of hemisegments with supernumerary Tin CBs.

Slit and its Robo receptors are known actors of CB migration, alignment and polarization^8,14,15,29^. Slit/Robo mediated cardiac morphogenesis is modulated by the transmembrane integrin receptors their ligands and intracellular interactors^12^. Taking into account the link between CAP and integrin adhesion complex and the alignment defects observed in *dCAP49e* embryos we tested whether Slit localization is affected in *CAP* and *MSP-300* mutants (Fig. 2 A’,B’,C’). We found a clear Slit localization defects in both mutant contexts. Slit was ectopically expressed in misaligned CBs creating a meshwork between CBs (Fig. 2B’,B”,C’,C”). Similar ectopic enrichment of Slit around mislocalized CBs has also been observed in cardiac specific knock-downs of CAP (miR strategy) and MSP-300 (short hairpin mediated strategy)^26^ (Fig. 2E, Sup. Fig. 2A-B’’). Collectively, these data show that *CAP* and *MSP-300* that accumulate in a dotty pattern ensure correct Slit localization in CBs. They also suggest a potential interaction between CAP, MSP-300 and integrin complex in migrating cardioblasts.

We then tested whether the cardioblast cell shape is affected in *CAP* mutants. Alpha-spectrin staining showed that *CAP-/-* cardioblasts are misarranged compared to the wild type where CBs are perfectly aligned along AP axis and matching with their contralateral partner (arrows and arrowheads in Fig 2D). In *CAP* hearts we observed changes in shapes and the orientation of CBs along the A-P axis (arrowhead in Fig. 2D’), misalignment leading to CBs matching defects (arrows and arrowheads, Fig. 2D-D’’) and mispositioning within rows of cells (star in Fig. 2D’) suggesting effected polarity and directionality of CBs migration. These defects occur also when CBs do not form clusters suggesting that clustering is only secondarily responsible of the observed phenotypes.

Finally, we tested genetic interaction between *CAP* and *MSP-300* using transheterozygous *dCAP49e/MSP-300^SZ75^* mutant embryos. In contrast to simple heterozygous controls, CBs alignment is affected in transheterozygous context (Fig. 2E, Sup. Fig. 2C-C’’), showing that *CAP* and *MSP-300* genetically interact and play convergent roles during CBs migration. We also tested whether changes in total cardiac cells number could account for *MSP-300* and *CAP* mutant phenotypes and found that knock out or cardiac specific KD of *MSP-300* and *CAP* both lead to an increase in the CBs number (Fig. 2F-K). Interestingly, only Tin-positive CBs population is enlarged suggesting a specific function of CAP and MSP-300 in cell cycle arrest in Tin-CBs (Fig. 2F-J, 2L, Sup. Fig. 2D), thought to undergo symmetric cell divisions.

### Vinculin-Beta Integrin complex is required for CAP and MSP-300 polarized localization

Cap protein interacts directly with Vinculin (Vinc) at the muscle attachment sites where it is recruited to the Integrin complex to modulate Actin cytoskeleton^26^. Therefore, we asked whether Vinc is also present in CBs during embryonic development and could interact with CAP. We took advantage of a recently generated Vinc-GFP knockin allele allowing to follow Vinc localization *in vivo*. We detected restricted Vinc-GFP signal at the apical and basal membranes of CBs reminiscent of that of CAP and MSP-300 (arrows Sup. Fig. 3A-B’’’) and recalling the localization of other members of the integrin adhesion complex such as βPS-integrin (arrows Sup. Fig. 3 C-C’’’) or Talin ^12,13^.

To test whether Vinc is required for CAP localization we analyzed the impact of heart-specific *Vinc* Knock Down on both CAP and MSP-300 expression. We observed that morphology of the heart proper is affected with inclusion of CBs (Fig. 3B, arrowhead) indicating their misalignment during migration. Dots of co-accumulation of CAP and MSP-300 are still present, however their pattern is disorganized suggesting that polarization of CBs is defective (arrows Fig. 3B-B”). Thus, during CBs migration dotty CAP accumulations in CBs depend on MSP-300 function while the involved in cell adhesion Vinculin is required for the polarized pattern of these large protein accumulations. Cardiac specific KD of Vinc also induces a meshwork of Slit (Fig.3 C-C’’) and an increase in Tin CBs number in some hemisegments reminiscent of CAP and MSP-300 loss of function (Fig. 3 D). Similar results are also obtained in cardiac specific KD of βPS-integrin (Sup. Fig. 3 D-D’’). Finally, we tested if the Integrin ligand Laminin A also known to interact genetically with *slit^21^* is affected in the loss of *MSP-300* which could provide evidence that LINC complex have an impact on the ECM content of CBs. Interestingly, we observed a disorganized pattern of LamA in *MSP-300* homozygous mutant embryos with breaks in the basal lamina suggesting the requirement of Drosophila Nesprin like proteins for ECM structure (Sup. Fig. 4A-C’).

**Figure 3:**
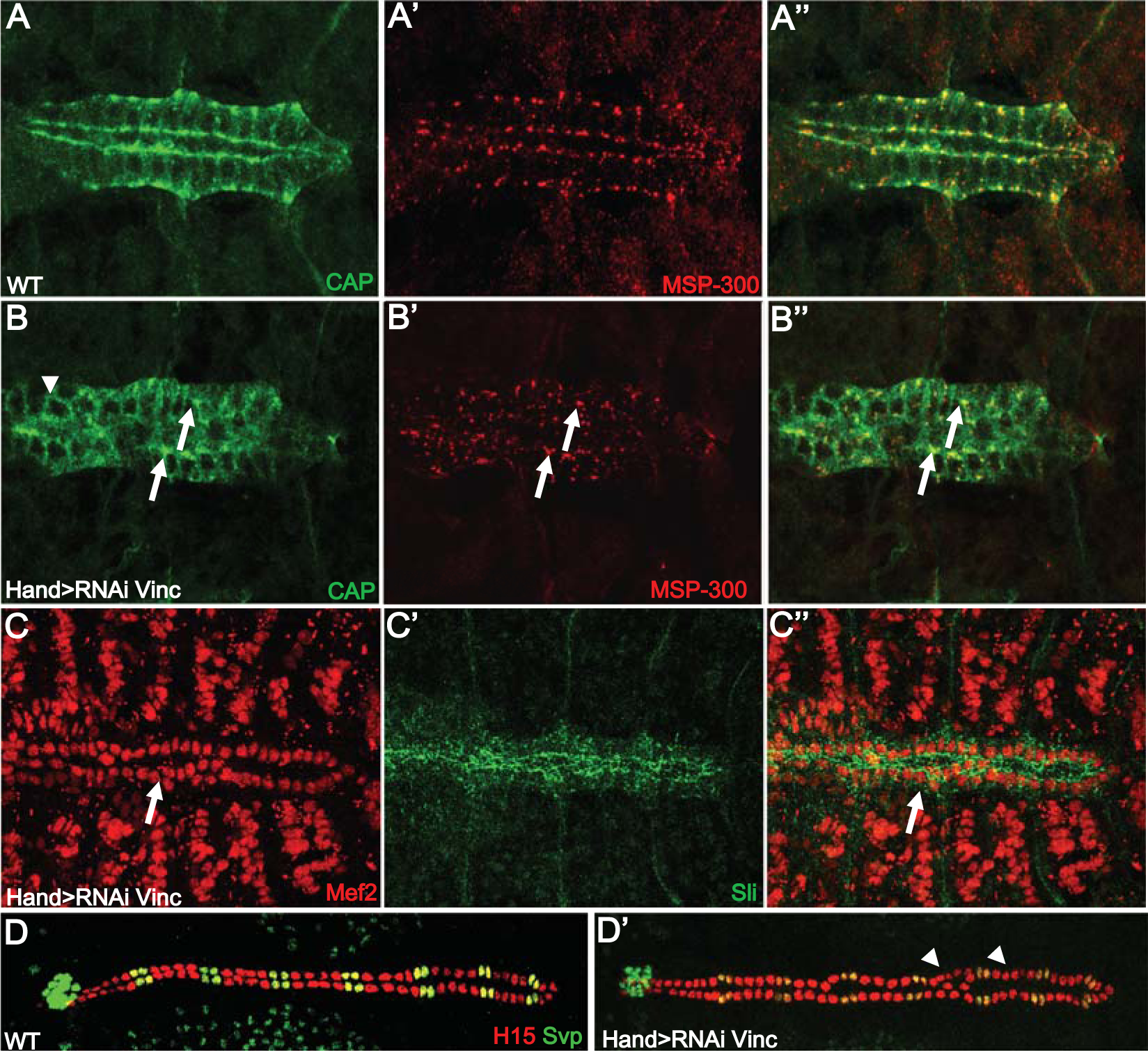
Vinculin controls polarized accumulation of CAP and MSP-300 in CBs. (A-B”) Immunostainings with anti-CAP and anti MSP-300 antibodies in WT and Cardiac specific *Vinculin* KD. Accumulation of CAP and MSP-300 is still visible but in a disorganized pattern suggesting polarity defects. (C-C”) Analysis of the effect of cardiac specific *Vinc* KD on CAP and MSP-300 localization. *Hand>RNAi-Vinc* leads to mislocation of CBs (arrowhead in B) and affects polarized localization of CAP and MSP-300 dots (arrows in B”). A disorganized pattern of CAP and MSP-300 is observed while dots seem to be maintained in *Vinc* KD. (D-D’) Compared to WT, *Hand>RNAi-Vinc* induces a specific increase of Tin CBs in 2 hemisegments (arrowheads)

All these observations reveal, that the polarized location of CAP-MSP-300-Vinc complexes play an important role in proper migration of CBs. One possibility is that they function by locally linking Vinc adhesion complexes to actin cytoskeleton. This would generate directional internal tensions that keep CBs aligned during their collective migration and could also regulate quiescent Tin-CBs state, cell shape changes during lumen formation and cardiac tube compaction.

### CAP and MSP-300 mediate F-actin bridging between nucleus and integrin complex

MSP-300 is the ortholog of mammalian Syne 1-2 and contains both actin binding and nuclear envelop localization domains while CAP binds to actin-binding proteins including Vinc. This suggests that F-actin might be a potential co-partner associated to the MSP-300-CAP-Vinc dots during CB migration. To test this hypothesis, we performed Phalloidin staining to reveal F-actin pattern in migrating CBs. Interestingly, F-actin accumulates on basal and apical sides of CBs very much like CAP and MSP-300 (Fig. 4 A-A’’’, Sup. Fig. 3 B-B’’’). Hence, we tested whether MSP-300 was required to stabilize F-actin accumulation at the interface of nucleus and CAP-Focal adhesion complex. Indeed, MSP-300 loss of function induced a strong decrease in localized F-actin expression in CBs (arrowheads Fig. 4 B-B’). Strikingly, this F-actin phenotype was specific for Tin positive CBs whereas Svp-CBs still accumulated F-actin in MSP-300 mutant background (arrows Fig. 4 B-B’) suggesting specific function of MSP-300 in Tin-CBs. We thus conclude that CAP and MSP-300 control coordinated CB migration and cell cycle exit by ensuring F-actin involving polarized link between nucleus and CB cell membrane. They exert their function essentially in “working CBs” population expressing Tin. F-actin stress fibers have been shown to exert mechanical tension on nucleus providing mechano-sensing properties to orient nucleus and modulate its shape during cell migration. Thus, we measured nucleus shape parameters in CAP and MSP-300 loss of function contexts to assess potential changes. We observed that comparing to the WT context, CAP and MSP-300 null mutants or cardiac specific CAP KD induce a significant decrease of nuclei volume and surface and an increase in flatness (Fig. 4C-E). These results show that upon loss of function of these factors nuclei are subject to deformation that could lead in fine to changes in gene expression.

**Figure 4:**
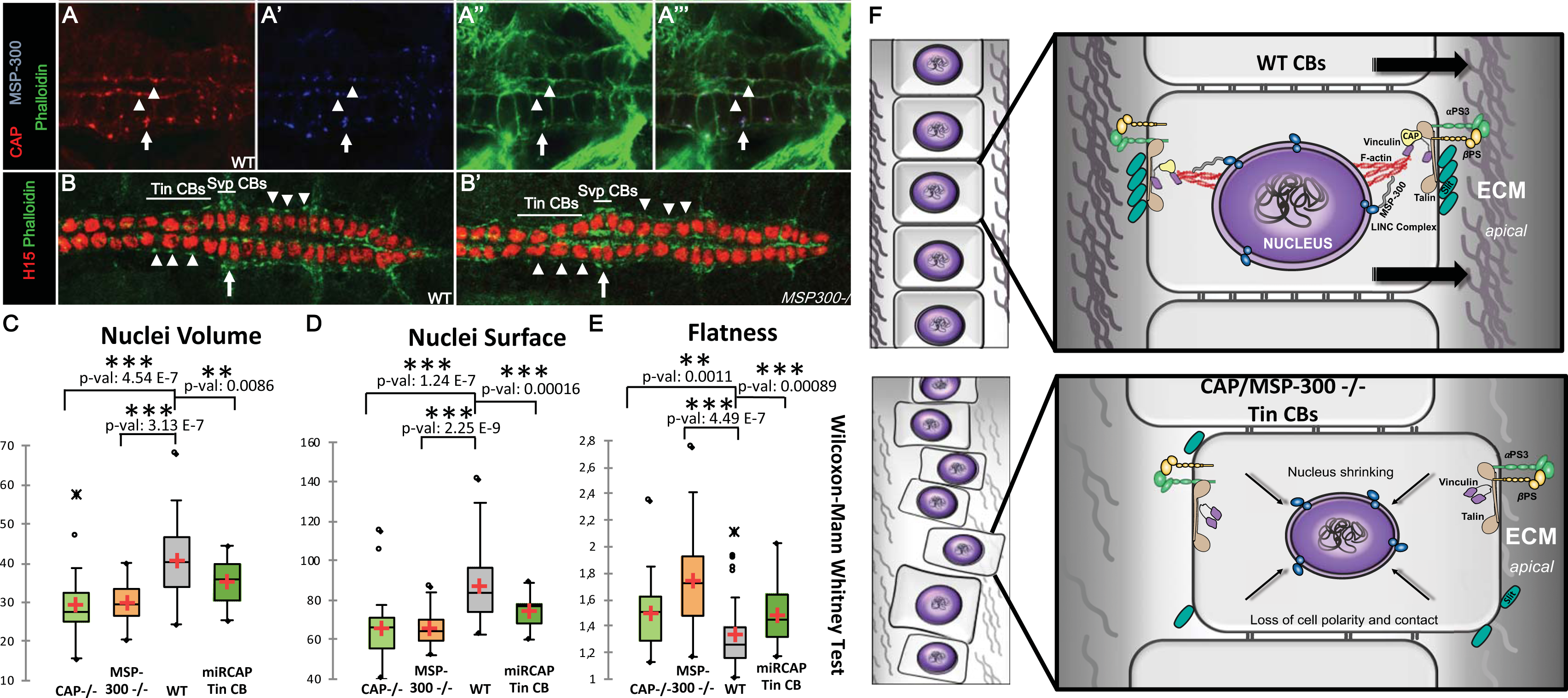
A polarized nucleus-cytoskeleton-ECM connection controls collective migration. (A-A”’) Dorsal view of heart proper showing colocalization of CAP, MSP-300 and F-actin on apical (arrowheads) and basal (arrows) sides of CBs showing a polarized link between nucleus and cell membrane on each side of the cells. (B-B’) Dorsal view of posterior part of the heart showing that loss of MSP-300 (mildly affected embryo) leads to a decrease in F-actin polymerization more particularly in Tin CBs (arrowheads). (C-E) Nuclei morphology parameters assessment using nucleus J software in different genetic context. (C) Nuclei volume is significantly reduced in *CAP*, *MSP-300* mutant CBs and in cardiac specific KD of *CAP*. (D) Nuclei Surface is also significantly reduced in the same contexts. (E) In contrary, flatness of CB nuclei increases in *CAP* and *MSP-300* mutant contexts. (Wilcoxon-Mann Whitney Test, n=32) (F) Model of nucleus-cytoskeleton-focal adhesion complex involving CAP adapter protein and Nesprin protein MSP-300 stabilizing nuclei morphology and polarized link to cell membrane on apical and basal sides. In the loss of *CAP* or *MSP-300* contexts, Vinculin interaction with F-Actin is lost, leading to a weakening of focal adhesion and affecting Slit polarization. These defects in turn induce a decrease in cell-cell contacts between neighboring CBs and misalignment within the migrating CB rows. Impact on nuclei morphology is also observed, leading potentially to gene expression changes.

## Discussion

We analyzed expression and function of two novel actors of cardiac morphogenesis encoding an Integrin complex adaptor protein CAP and a Nesprin protein belonging to the LINC complex, MSP-300. Our study highlights a crucial role for these proteins during the late phase of CBs migration and reveals a connection between nucleus and extracellular matrix via integrin/vinculin adhesion complex required for a coordinated and directional cells movement (Fig. 4F). We also provide evidence that this nucleus-cytoskeleton-cell membrane link is important to maintain CBs polarity and contact between ipsilateral and contralateral CBs and for nucleus shape properties. It might also participate in locking cell divisions processes during the migration progression of Tin cardiomyocytes.

### CAP and MSP-300 are required for coordinated CBs migration

Among the genes identified by TRAP-seq involved in "cell migration" GO category we focused on *CAP* and *MSP-300*. Our data reveals that these two proteins colocalize in a very atypical dotty pattern under the basal and apical membrane of CBs from early stage 15 and until stage 16 during the final step of migration. Interestingly, Slit transiently accumulates at the same locations before being polarized on the apical CBs area where it is required for lumen formation. Functional analyses on *CAP* and *MSP-300* lead us to the conclusion that they act together to ensure the correct alignment of CBs during collective migration and Tin CBs cell cycle arrest. These mechanisms seem only partially coupled as we observed wrong alignments of CBs even when the two rows are present as a bilayer without CB clusters. Slit localization is also affected in the mutants of these two genes suggesting that Slit mediated-directionality of CBs migration might be affected. Indeed, a conserved aspect of Slit/Robo signaling is its involvement in cell guidance mechanisms^16,30,31^. In *Drosophila*, Slit is known to interact genetically with several members of the integrin complex (αPS3, βPS) and integrins are required for apical localization of Slit and leading edge activity. Integrin-based adhesion complexes are crucial for cell migration. The direct coupling between the cell and the ECM at integrin-based adhesion sites allows cells to mechanically sense their physical surroundings and adjust mechanisms of migration by initiating downstream signaling events that regulate cytoskeletal organization. In *Drosophila* CAP has been described as a scaffolding protein at membrane-cytoskeleton interfaces required to facilitate the assembly of protein complexes involved in cytoskeletal regulation and modulating membrane stiffness. Furthermore, a direct interaction between CAP and Vinculin has been shown at muscle attachment sites^26^. Vinculin is a key player in focal adhesion where it interacts directly with Talin and ensures a link between adhesion proteins and actin cytoskeleton. Recent data also suggest that Vinculin could regulate directionality and polarity during cell migration^32,33^.

In CBs, loss of Vinculin leads to disorganized CAP and MSP-300 dotty localization, probably due to the loss of Vinc-anchoring points at the cell membrane, resulting in affected CBs migration.

MSP-300 belongs to Nesprin family of protein and possesses several protein isoforms containing different lengths of spectrin repeats allowing interaction with actin and a nuclear SUN protein interaction domain KASH. MSP-300 is a member of the LINC complex linking actin cytoskeleton to the nucleus. LINC complexes are known to be important in transmitting mechanical stresses from the cytoskeleton to the inside of the nucleus during many cell processes such as migration. This mechanical nuclear-cytoskeleton coupling combined with adhesion to the external environment mediates the transduction of forces to the nucleus that might be involved in controlling directional and synchronous cell movements as well as gene activities. Our results support this view. We propose that MSP-300-CAP interaction ensures the connection of nuclei to adhesion complex via actin cytoskeleton and Vinculin and could thus modulate guidance signaling and CBs leading edge dynamics. Further analyses will be required to distinguish the specific functions of these two proteins in CBs migration as well as between CB cell types.

One of the remaining questions relays on the fact that CAP-MSP-300 dots accumulate at both sides of migrating CBs. Recent evidences suggest the existence of an actin structure called perinuclear actin-cap that interact with specific actin-cap-associated focal adhesion components at two positions in the direction of the aligned actin-cap fibers, at leading lamellipodial edge and the trailing edge of the cell. This newly identified actin structure is linked to the nucleus via the LINC complex^34,35^ and might be necessary to coordinate cell movement and nucleus shape during migration. Further work will determine if this perinuclear actin-cap plays indeed a role in CBs migration during *Drosophila* embryogenesis.

Transition from cell cycle exit to cell migration is a coordinated process that requires multilevel controls in terms of cell polarity, cell adhesion and differentiation involving changes in gene expression programs. Several studies have made links between mechanotransduction mediated by focal adhesion or the LINC complex and cell cycle control. Our results provide evidence of a potential impact of these interconnected units to control cell divisions during CBs migration. Recent findings have shown that deleting components of the LINC complex abolished control of endoreplication in myofibers nuclei by affecting the chromatin regulator BAF leading to an increase DNA content^36^. Mutations in *nesprin-1* have been shown to reduce levels of Lamin A/C^37^, a known partner of BAF^38^, leading to changes in chromatin structure and aberrant expression of genes involved in differentiation^39^. We also observed a loss of Lamin C in CBs upon *MSP-300* loss of function (data not shown). Furthermore, these *nesprin-1* mutations lead to increased activation of the ERK pathway in mouse heart tissue^37^. In *Drosophila*, late EGFR mediated ERK activation has been shown to promote Tin CBs subpopulation^40^. According to our data, one hypothesis is that CAP and MSP-300 would counteract EGF pathway maintaining the correct number of Tin migrating CBs. Further investigations will be required to clarify the involvement of CAP and MSP-300 in these processes.

## Acknowledgements

This work was supported by Agence Nationale de la Recherche JC (CARDIAC-SPE) and FEDER/Region Auvergne Young Researcher grant. We thank Sophie Desset, Tristan Dubos and Caroline Vachias (GReD institute) for their technical assistance as well as CLIC facility.

## Experimental procedures

### *Drosophila* genetics

Drosophila *CAP* deletion mutant *dCAP49e* deletes 2.9kb downstream of CAP^CA06924^ P-element. The *dCAP49e* mutant and *UAS-CAPmiR* construct has been generated according to ^26^. *dCAP49e, P(UAS-CAP.miRNA)3* were from Bloomington Stock Center. *MSP-300*^*SZ75*^ mutant is a gift from Talila Volk (Weizmann Institute of Science, Rehovot, Israel). It carries truncation of the C-terminal part of Msp-300^41^. MSP-300 TRIP lines (32377, 32848) and myospheroid TRIP lines (27735, 33642) from Bloomington Stock Center. Vinculin RNAi lines (v34585, v34586) were obtained from the Vienna Drosophila Research Center and TRIP line (25965) from Bloomington Stock Center. Vinculin-GFP line is tagged at its N-terminus with superfolder GFP using a pFlyFos025866 Fosmid which encompasses the 8 kb of Vinculin gene along with 23.4 kb upstream and 6.8 kb downstream regions integrated in attp2 site^42^.

### Immunostaining

*Drosophila* embryos were processed for immunostaining as described previously for most of the experiments ^43^. Heat fixation were also performed on WT dechorionated embryos for 10 second in boiling water followed by 5 min devitellinization in a 1:1 mix of heptan and methanol on shaker. The following antibodies were used: Rabbit anti-CAP 1:1000 (a gift from A.L Kolodkin, Johns Hopkins University School of Medicine, Baltimore, USA), Rabbit anti-Mef2 1:1000 (a gift from M.V Taylor, School of Biosciences, Cardiff University, Cardiff, UK), Guinea Pig anti-MSP-300 1:2000 (a gift from T. Volk, Weizmann Institute of Science, Rehovot, Israel), Guinea Pig anti-H15 1:2000 (a gift from J.B Skeath Washington University School of Medicine, St. Louis, USA), mouse anti-Slit at 1:100 (DHSB), anti-Vinculin at 1:50 (DHSB), anti-Seven up at 1:100 (DHSB), anti-Armadillo at 1:100 (DHSB), anti α-Spectrin 1:100 (DHSB), Alexa fluor 488 Phalloidin (Invitrogen A12379).

### Optical cross section of Drosophila embryonic heart tubes

IMARIS software 7.6.5 has been used to open images in a 2D display mode (Slice view).

In the menu bar we selected “Edit – Crop 3D”. A rectangle, with yellow borders, will overlay on all three views. It represents the region of interest and it can be modified in size and position resulting in the cross-sections of the selected part of the heart tube.

The Surpass mode is used to create a 3D reconstruction of the *Drosophila* embryonic heart tubes, and setting the pointer on Navigate mode allows to rotate the resulting object through a range of angles. This was done in two different regions: aorta and the heart proper.

### Nuclei morphology measurement

Microscopic observations were performed by confocal microscopy to produce images using Leica SP8. All images were acquired using a 40× oil objective with a numerical aperture of 1.3, a pixel size of 0.054 µm (x,y) and voxel depth of 0.2 µm. Furthermore, all initial anisotropic voxels are converted to isotropic voxel (i.e. cubic, xyz=0.1 µm) prior to calculation^44^. The ImageJ plugin NucleusJ was used to characterise nuclear morphology. A detailed description of the quantitative parameters generated by NucleusJ can be found in supplemental materials of Poulet et al.^44^.

