## Supplementary Figures for "A polarized nucleus-cytoskeleton-ECM connection controls collective migration and cardioblasts number in *Drosophila*"

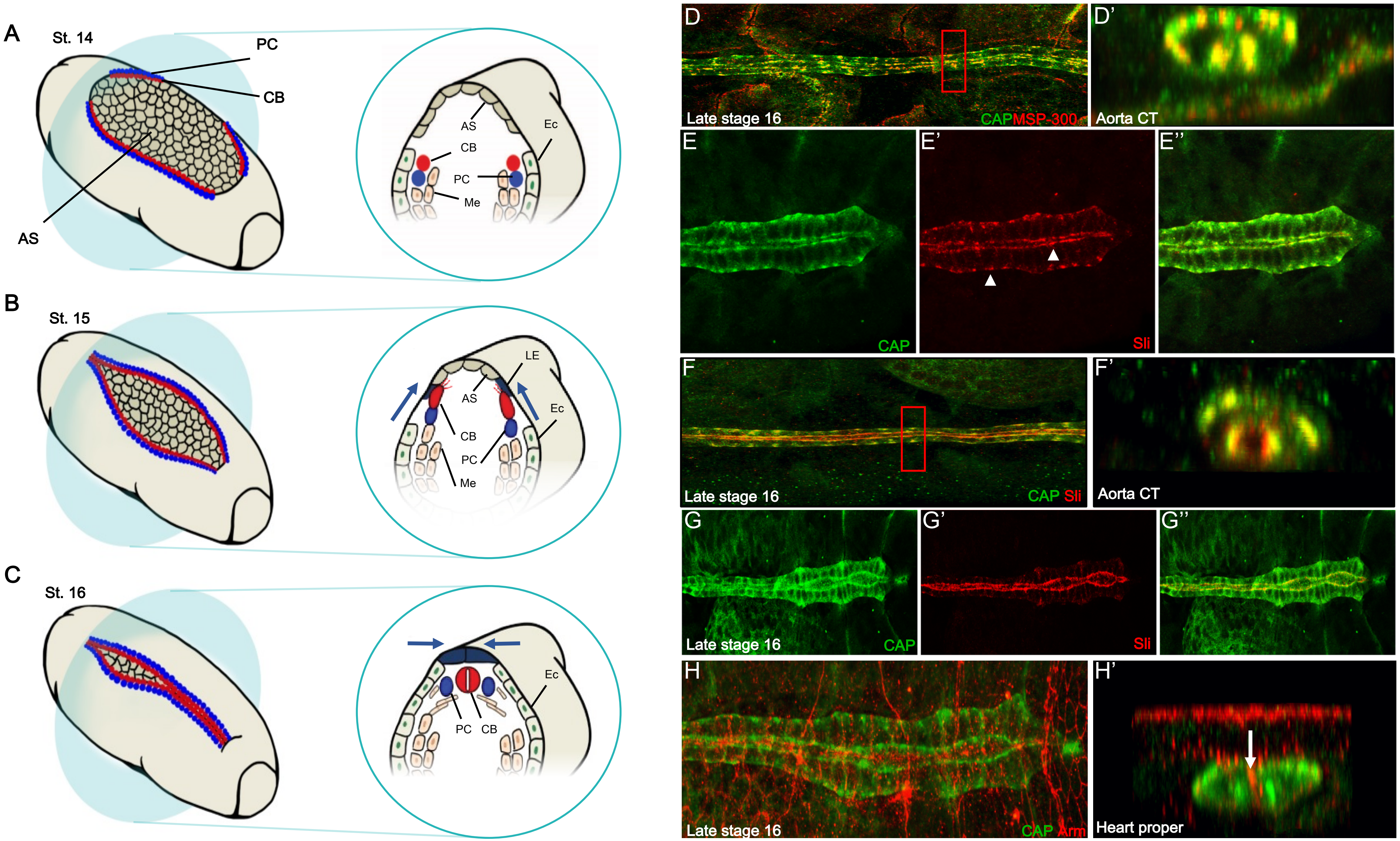


**Figure supplemental 1: CAP expression pattern in cardiac cells**

(A-C) Schematic representation of heart closure processes that have been targeted for this study. At Stage 14, CBs migrate coordinately with ectoderm during dorsal closure. By stage 15 autonomous movements through filopodia present at the leading edge are controlling CBs migration. At stage 16, the two rows of CBs contact each other and the process of lumen formation begins.

(D-D') Dorsal view of a stage 16 embryo showing CAP and MSP-300 localization in aorta and 3D reconstruction of transversal cut in aorta validating accumulation of MSP-300 and CAP on basal and apical side of CBs.

**Figure supplemental 1: CAP expression pattern in cardiac cells (next)**

(E-E’’) Dorsal view of a stage 16 embryo showing that Slit colocalizes with CAP protein in a similar dotty pattern on apical and basal sides of CBs (arrowheads).

(F-F') Dorsal view of a stage 16 embryo showing CAP and Slit localization in aorta and 3D reconstruction of transversal cut in aorta validating accumulation of Slit and CAP on basal and apical side of CBs. Note that Slit protein is secreted at luminal side explaining the partial overlap with CAP.

(G-G’’) At late stage 16, Slit protein becomes accumulated at the lumen and strongly reduced at the basal side (arrows), while CAP is still maintained on both sides.

(H-H’) CAP is not colocalized with Armadillo at the adherens junctions between contralateral CBs (arrow).

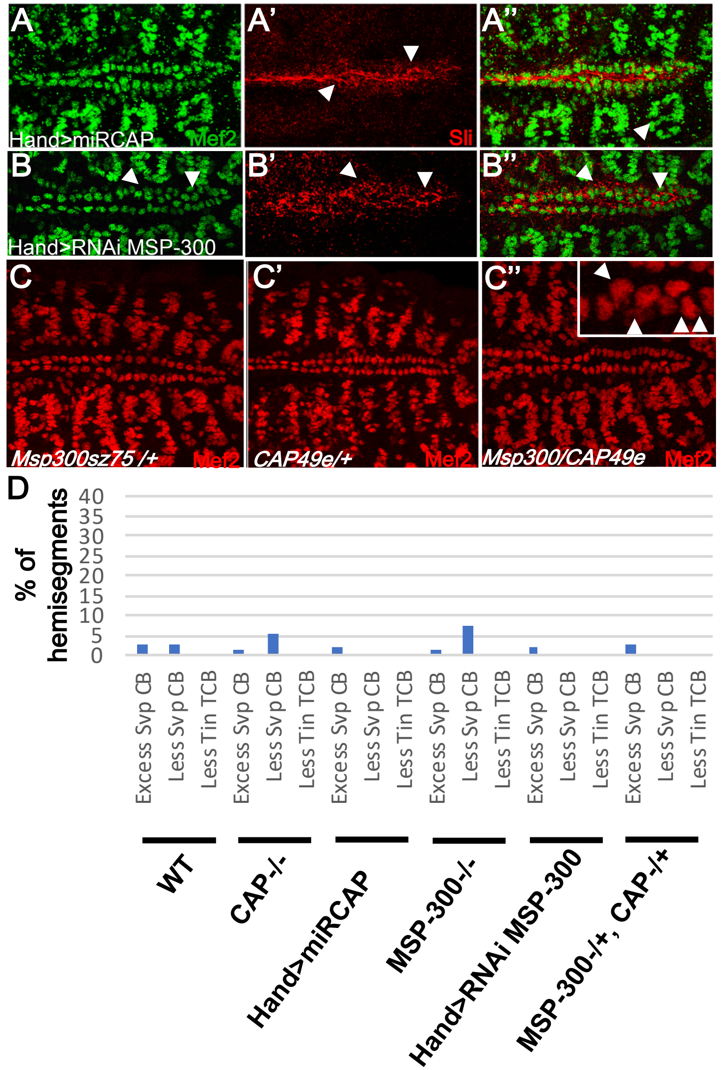


**Figure supplemental 2: CAP and MSP-300 interact genetically to control CBs alignment and number**

(A-A") Cardiac specific miR KD of *CAP* using Hand-Gal4 driver induces CBs misalignment and polarity defects visualized with anti-Mef2 and anti-Slit antibodies (arrows) similarly to *CAP* homozygous mutant.

(B-B'') Cardiac specific short hairpin KD (TRIP line) of *MSP-300* using Hand-Gal4 driver induces CBs misalignment and polarity defects visualized with anti-Mef2 and anti-Slit antibodies (arrows) similarly to *MSP-300* homozygous mutant.

(C-C'') Immunostaining with anti-Mef2 in *CAP* and *MSP-300* single heterozygous (C-C') by comparison with double heterozygous contexts (C''). No apparent change in CBs alignment can be observed in single heterozygous background while clusters of CBs in heart proper are visible in transheterozygous context (C'', arrowheads in higher magnification window) demonstrating a genetic interaction between *CAP* and *MSP-300* during CBs migration.

(D) Histogram showing the percentage of hemisegments with changes in CBs number other than an excess of Tin CBs.

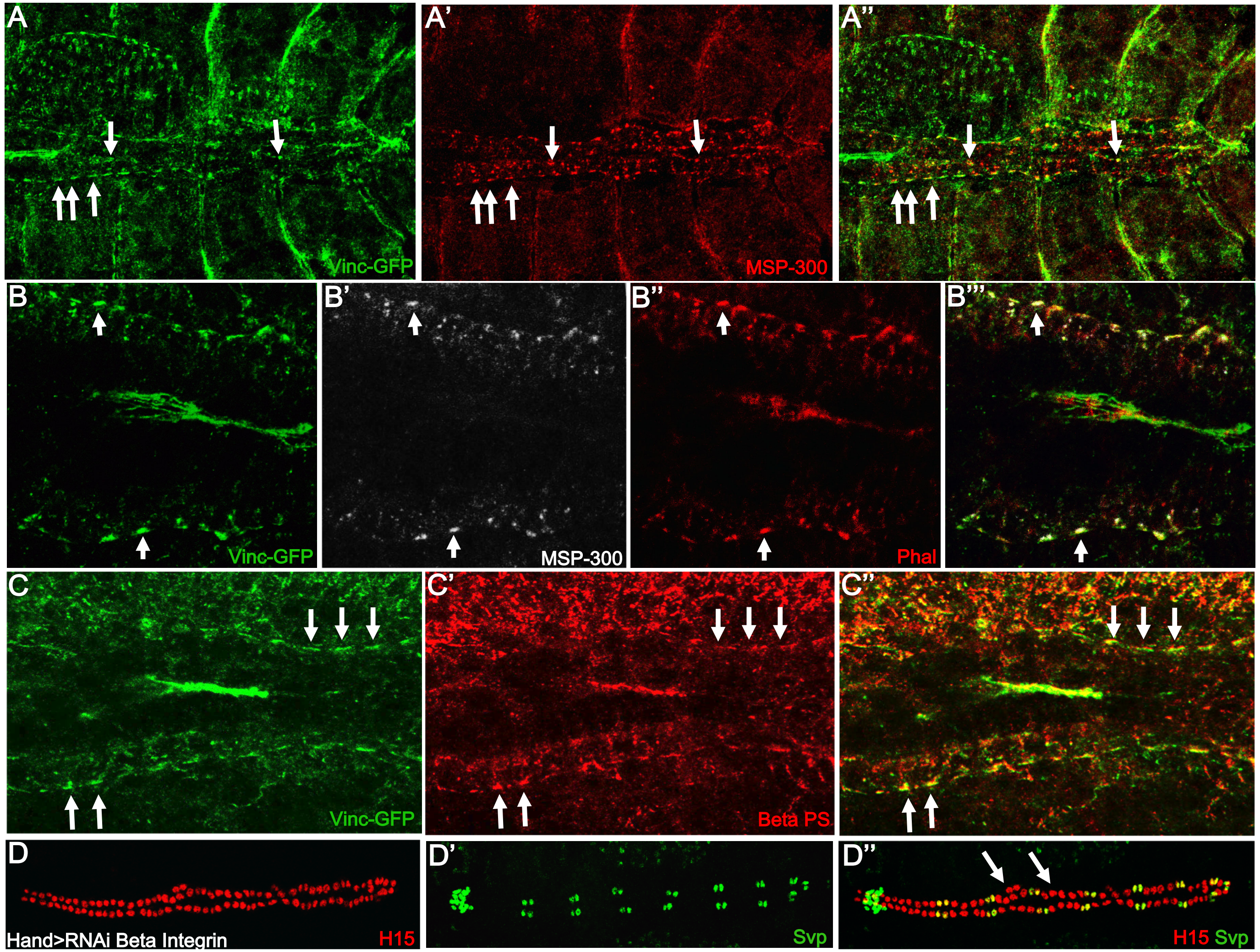


**Figure supplemental 3: Vinculin expression in cardioblasts**

(A-A'') Dorsal view of stage 16 embryos revealing Vinculin-GFP localization in a spotty pattern in close proximity to MSP-300 accumulation on basal and apical sides (arrows).

(B-B'') Higher magnification pictures showing that MSP-300 is accumulated where the maximum of Vinc-GFP is present. Vinculin partner F-actin is also present in a similar dotty pattern (B"-B"')

(C-C") Vinc-GFP colocalizes with βPS Integrin in cardioblasts (arrows) suggesting a focal adhesion based process connected to the nucleus through F-Actin.

(D-D") Cardiac cells inducible knock down of βPS Integrin with Hand-GAL4 driver leads to an increase in Tin CBs in some hemisegments (arrows in D").

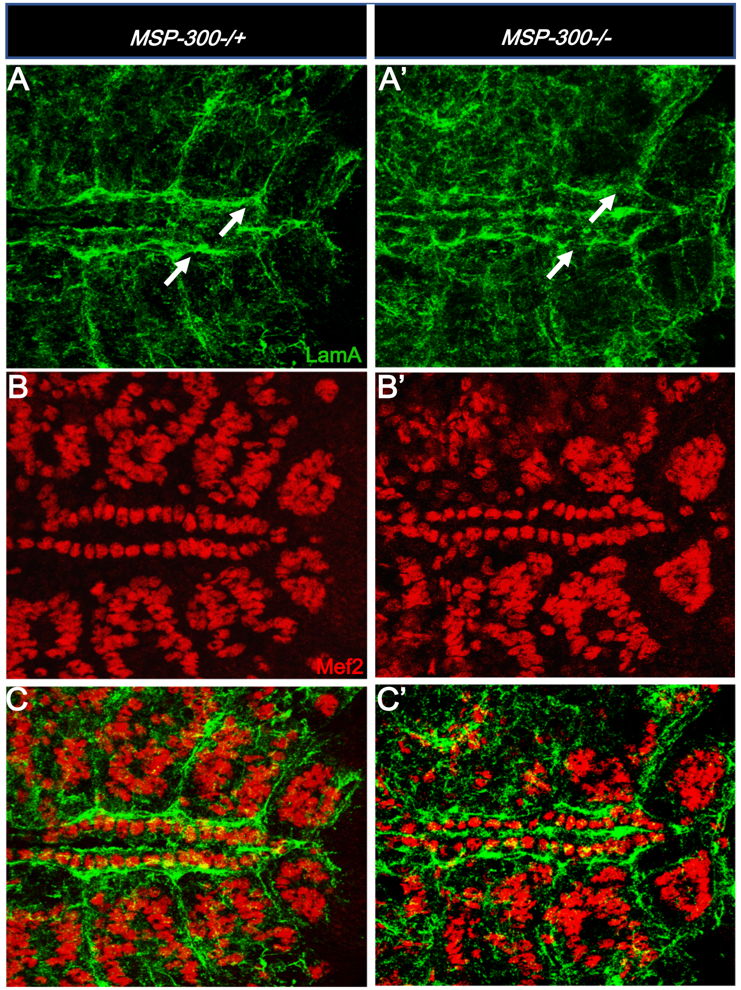


**Figure supplemental 4: ECM protein Laminin A localization is affected in *MSP-300sz75* mutant context**

(A-C') Immunostainings showing that loss of MSP-300 induces a disorganized pattern of Laminin A on basal and apical sides of CBs. LamA basal layer is thinner and holes are visible around some CBs (arrows in A-A').
